# Phase resetting in human stem cell derived cardiomyocytes explains complex cardiac arrhythmias

**DOI:** 10.1101/2025.09.24.678279

**Authors:** Khady Diagne, Thomas M. Bury, Morgan E. Pettebone, Marc W. Deyell, Zachary Laksman, Alvin Shrier, Leon Glass, Gil Bub, Emilia Entcheva

**Author notes:** These authors contributed equally to this work (Khady Diagne, Thomas M. Bury). These authors contributed equally to this work (Gil Bub, Emilia Entcheva).

## Abstract

Phase resetting of cardiac oscillators underlies some complex arrhythmias. Here we use optogenetic stimulation to construct phase response curves (PRC) for spheroids of human induced pluripotent stem cell derived cardiomyocytes (iPSC-CM) and a computational cardiomyocyte model to identify ionic mechanisms shaping the PRC. The clinical utility of the human PRCs is demonstrated by adding a patient-based conduction delay to the same equations to explain complex multi-day Holter ECG dynamics and cardiac arrhythmias. Periodic stimulation of these patient-based models and the computational model of human iPSC-CM reveal similar bifurcation patterns and entrainment zones. Cell therapy by injecting iPSC-CM into diseased hearts can induce ectopic foci-based engraftment arrhythmias. The PRC analysis offers a potential strategy to entrain these foci in a parameter space that avoids such arrhythmias.

---

From the first (*1*) to the last (*2*) heartbeat, biological systems display sudden dynamic transitions that can be analyzed mathematically (*3*). Such transitions can be elicited by periodic stimulation of spontaneous oscillators. For example, perturbing the frequency of periodically beating heart cells leads to a panoply of simple and complex rhythms (*4*). Phase response curves (PRCs) (*4-6*) are compact representations of the responsiveness of cardiac regions with innate oscillatory dynamics to small perturbations from an external driver and can predict patterns of synchronization. PRCs have been useful in analyzing the complex dynamics of coupled biological oscillators (*7*). An example is parasystole – an arrhythmia that arises when a cardiac region with automaticity surrounded by tissue with decreased conduction competes with the native pacemaker (the sinus node) (*6, 8-11*). In parasystole, the abnormal focus is the source for premature ventricular complexes (PVCs). Despite extensive research on parasystole, there are major gaps in the translation of this understanding to the clinic. This knowledge is also relevant to optimize an emerging therapy for cardiac regeneration involving the transplantation of human induced pluripotent stem-cell derived cardiomyocytes (hiPSC-CM) into a diseased heart, an approach that has moved to clinical trials (*12-14*). From a dynamics perspective, the engraftment of spontaneously beating hiPSC-CM into the heart closely resembles parasystole. A major complication associated with this therapy is the occurrence of potentially fatal cardiac rhythms, termed engraftment arrhythmias. These have been observed in large animal models within the first two weeks post-transplantation (injection) (*14-17*). Electrocardiographic records from such studies show that at least in part, engraftment arrhythmias may result from competing oscillators (the sinus node and the transplantation site) (*17-19*).

In previous work, the dynamical responses of spontaneously beating cardiac tissue from embryonic chicks (*4*) and dogs (*20*) to external periodic stimulation have been represented by finite difference equations, describing phase resetting. Here, we sought to extend this PRC-based analysis to capture the dynamics of hiPSC-CM using experimental and computational approaches to better understand the potential role of phase resetting in the generation of PVCs in patients following stem cell grafts as well as in patients with frequent PVCs. Using an ionic model of these hiPSC-CM, we suggest a strategy to decrease the prevalence of ectopic activity.

## Phase response curves for hiPSC-CM spheroids

To obtain a PRC of human-derived tissue, we used self-assembled spontaneously beating spheroids of hiPSC-CM paired with light-responsive spheroids of non-beating human embryonic kidney cells to form a tandem cell unit (*21*), Fig. 1A. We stimulated the light-responsive spheroid with periodic depolarizing light pulses at well-defined frequencies which excited the hiPSC-CM spheroids (see materials and methods). Electrical responses (action potentials, APs) evoked in the hiPSC-CM spheroids were measured optically using a voltage-sensitive dye (*22-24*).

**Fig. 1:**
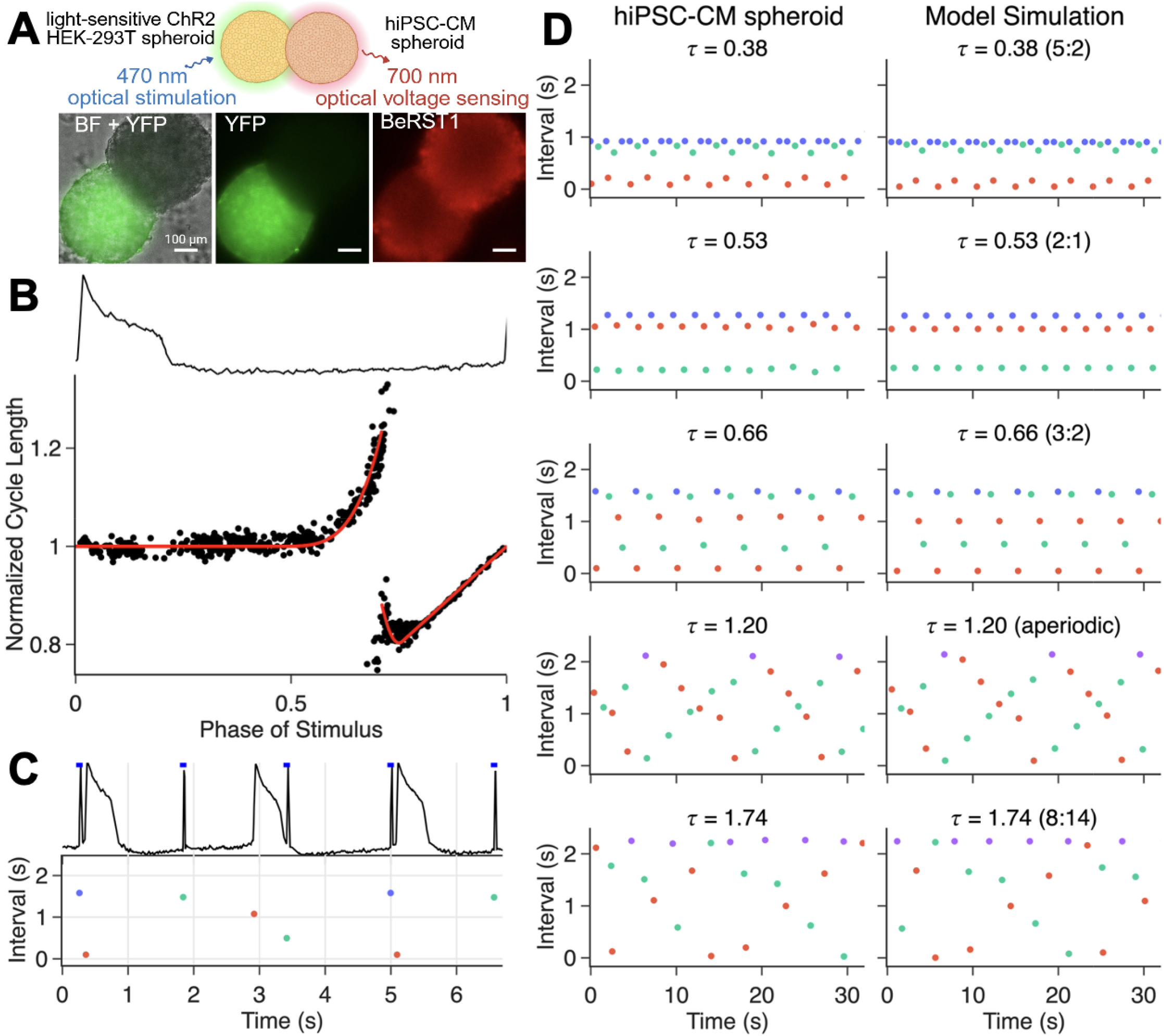
Phase response curve for human iPSC-CMs spheroids. (**A**) All-optical interrogation of cell spheroid pairs, shown in brightfield (BF), YFP fluorescence (YFP is a reporter tag for the optogenetic actuator ChR2), and BeRST1 (optical voltage sensor). (**B**) Trace of a full action potential cycle (top) and the phase-response curve (bottom), showing normalized cycle length as a function of the phase of the stimulus within the intrinsic cycle (2.4 s). Black dots show experimental data; the red line is the best fit of Eq. 2 (A = 60, = 0.71, B = 50, S= 0.80, RMSE=0.056). (**C**) Dot plot representation of the pacing data. stimulus-stimulus (blue); stimulus-AP (red); AP-stimulus (green); AP-AP (purple, not shown). (**D**) Entrainment patterns observed for various pacing frequencies in the experiment (left) and the periodically-forced oscillator model (right). is defined as he ratio of the stimulus cycle length to the intrinsic cycle length of the hiPSC-CMs. For periodic rhythms, locking ratios are given in parentheses in the form N:M, denoting N stimuli per M cycles of the forced oscillator.

Let *T* denote the spontaneous cycle length of the spheroid and *τ* = *T* _stim_/*T* be the normalized pacing cycle length, where *T*_Stim_ is the period of external stimulation. The light stimuli were 10 ms and delivered at multiple values of *τ*. For each stimulus *i*, delivered at phase *ϕ*_*i*_, we measure the change in cycle length in the hiPSC-CM spheroid. Sweeping over multiple stimulation frequencies allows the full range of phases to be sampled, generating the PRC for the hiPSC-CM spheroid (Fig. 1B). The resultant dynamics are shown in Fig. 1C by plotting the intervals between consecutive events using colored dots. We fit a piecewise-defined polynomial equation *g*(*ϕ*) to this experimental PRC (Fig. 1B, see materials and methods). The spread of points around this red curve (RMSE = 0.056) reflects small long-term effects of the stimuli, measurement noise, and intrinsic biological noise. We use this equation to model the hiPSC-CM spheroid’s response to periodic stimulation (*25, 26*). Let *ϕ*_*i*_ be the phase of the 11-th stimulus in the oscillator’s cycle. Then the phase of the next stimulus is given by

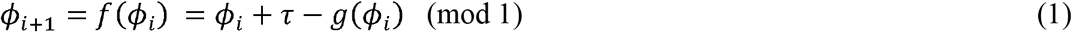

Simulation of this model corroborates the experimental observations, Fig. 1D. The dynamics are periodic with period N if *ϕ*_*i*_ = *ϕ*_*i* +*N*_ and *ϕ*_*i*+*j*_ ≠ *ϕ* for 0 < *j* < *N*; N:M periodic rhythms represent a repeating sequence of N stimuli and M action potentials. These experimental findings are consistent with the behavior of a perturbed oscillator (Eq. 1), with a strongly attracting limit cycle, i.e. following each stimulus, the cycle is reestablished, perhaps with a shift of phase.

Similar PRCs were obtained from six different stimulated hiPSC-CM spheroids (fig. S1, table S1). These human cardiac PRCs have: (i) A well-defined discontinuity separating cycle prolongation from shortening; (ii) Stimuli delivered early in the cycle have negligible effect on the cycle length or increase it; (iii) Stimuli delivered after the discontinuity lead to cycle shortening, with a “hook” followed by a distinctly linear segment. These features are consistent with previously reported experimental (*4, 20*) and theoretical (*27, 28*) studies of PRCs of non-human cardiac oscillators. The “hook” in the human cardiac PRCs obtained here, immediately past the discontinuity, was more pronounced than in previous studies from different species. Near the discontinuity (phase *ϕ*_*r*_), a stimulus either delays or triggers the next beat. This may reflect the influence of stochastic fluctuations due to ion channel behavior (*27*) or due to unrelated noise.

### Modelling dynamics of PVCs in clinical records using the experimentally derived PRC

Multi-day Holter recordings from 53 patients with more than 5% PVCs enrolled in the British Columbia PVC Registry were used. We developed a model of modulated parasystole incorporating the experimentally derived PRC and data-driven techniques to fit the model to individual patient ECG recordings (see materials and methods). The empirical PRC helped constrain the number of free parameters in the model in a biologically-grounded manner.

The clinical data was initially screened to select patients in whom more than 80% of the PVCs had the same morphology. This criterion yielded a subset of 40 patients. Identification of cycling coupling intervals — a hallmark of parasystolic dynamics (*29*)– further selected 7 out of the 40 patients (figs. S2-S3). For these patients, we investigated the extent to which the modulated parasystole model could reproduce the clinically observed dynamics.

The modulated parasystole model builds on earlier work (*11*) by including a conduction delay into and out of the ectopic focus (*30*). The model assumes a sinus pacemaker with period ⍰ _⍰_, an ectopic pacemaker with period 11, a refractory period ⍰, and modulation of the ectopic pacemaker by sinus beats via a PRC (⍰). As observed clinically, the model assumes that sinus beats immediately following an ectopic beat are blocked, and *vice versa*. Let ⍰ _⍰_ be the phase of the ⍰ ^th^ sinus beat in the ectopic cycle, determined by the timing of the sinus beat relative to the previous ectopic beat (Fig. 2A). Then the subsequent phase is given by:

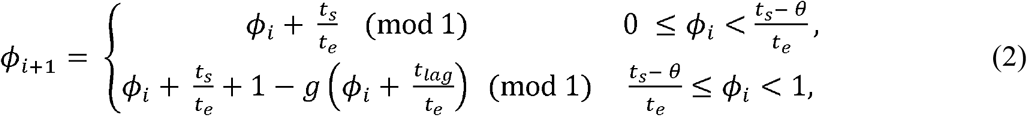

where ⍰ _lag_ is the total conduction delay into and out of the ectopic focus, and other parameters are as previouslydefined. The shifted phase ⍰ + ⍰ _lag_/⍰ _⍰_ represents the phase at which the sinus beat stimulus arrives at the ectopic focus (fig. S4). Accounting for this conduction delay is necessary to reproduce the dynamics observed in clinical recordings.

**Fig. 2:**
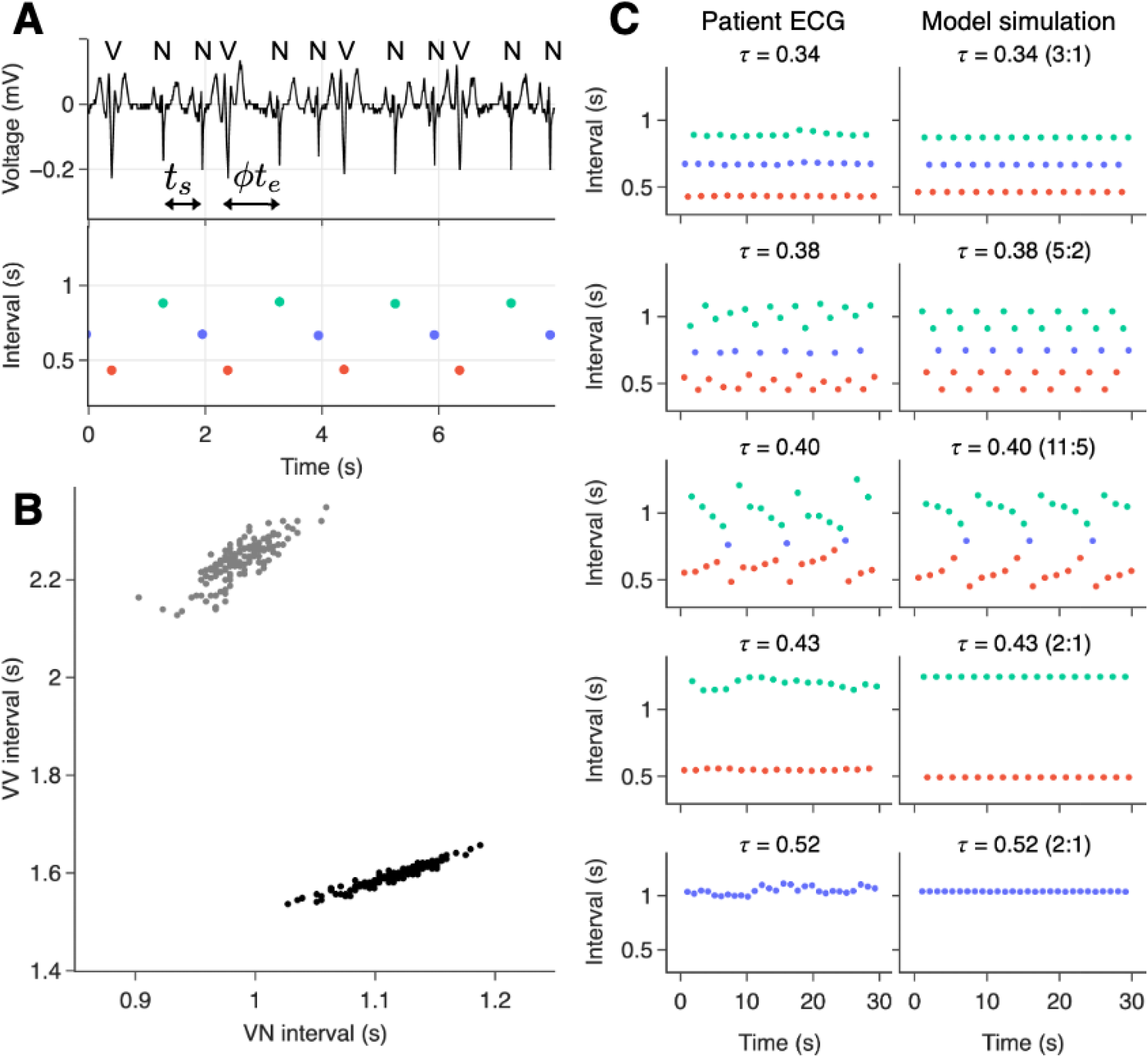
PVC dynamics in a patient with frequent PVCs and corresponding simulations of the modulated parasystole mode. **(A)** A segment of the patient’s electrocardiogram (top) and associated inter-beat intervals (bottom). N denotes a sinus beat, V an ectopic beat (premature ventricular complex, PVC). The sinus cycle length is denoted by t_s_, the ectopic cycle length by t_e_ and □ is the phase of the sinus beat in the ectopic cycle. Intervals are color-coded: NN (blue), NV (red), and VN (green). **(B)** A representation of the phase response curve of the ectopic focus, inferred from sequences of bigeminy (alternating sinus and ectopic beats, black dots) and trigeminy (two sinus beats between ectopic beats, gray dots) (see Methods). **(C)** Patient ECG recordings (left) and simulations from the modulated parasystole model (right), shown at five different ratios between sinus and ectopic cycle lengths (). The locking ratio of the model is given in parentheses in the form N:M, denoting N sinus cycles per M ectopic cycles.

The model includes 10 parameters (table S2), which are fit to each patient (see materials and methods for fitting procedure). We can reconstruct part of the hypothesized ectopic pacemaker’s PRC using the ECG (Fig. 2B, fig. S5) (*31*). For the model, however, *g*(*ϕ*) is obtained from the experimentally derived PRC which spans the entire oscillator’s cycle. Three parameters are estimated directly from the clinical data: the sinus period ⍰ _⍰_, the refractory period ⍰, and the slope of the PRC after the discontinuity ⍰ (figs. S6-S7). The remaining three parameters: ectopic cycle length ⍰ _⍰_, conduction delay ⍰ _lag_, and PRC discontinuity ⍰ _⍰_, are chosen to minimize the mean absolute error between the model output and the clinical recordings. Table S3 shows the unique set of fitted parameters for each patient.

In Fig. 2C we show dynamics from a single patient alongside model output using the best-fit parameters for five representative values of ⍰ _⍰_. The model’s ability to reproduce dynamics over the entire recording was assessed using the Kolmogorov–Smirnov test, i.e. a good fit (⍰ > 0.05) was considered when the inter-beat interval distribution from the simulation was not significantly different from that of the patient record. Allowing for a ±5% variation in the fitted parameters, the proportion of the recording that was well reproduced by the model ranged from 63.4% to 95.0% across patients (table S3).

### Empirical bifurcation analysis

We use the empirical PRC in Fig. 1B to simulate the dynamics in a periodically stimulated hiPSC-CM spheroid (Eq.1) and patient ECGs with recurrent PVCs (Eq. 2). We perform a bifurcation analysis of these models to explore the dynamical landscape of these systems (Fig. 3). From iteration of Eq. 1, we compute the bifurcation diagram, i.e. the long-term stimulus phases as a function of ⍰ for 0 ≤ ⍰ ≤ 1.8 (Fig. 3A). The diagram reveals a variety of stable periodic behaviors, interspersed with chaotic regions. A good agreement is seen with the experiments (overlayed in blue), despite noise. In Fig. 3B, we plot the experimental results from Fig. 1C in a return map of the phase of the stimulus in the oscillator’s cycle length in addition to the model’s iterations. This comparison allows us to distinguish between noisy periodic dynamics (⍰ = [0.38, 0.53, 0.66, 1.74]) and chaotic rhythms (⍰ = 1.2). The shaded area in the bifurcation diagram of Fig. 3A highlights the range of ⍰ values seen in the patient. We plot in Fig. 3C the bifurcation diagram for the parasystole model in Eq. 2 for 0.3 ≤ 11 ≤ 0.55, which follows very similar bifurcations as those seen in the simpler model above. In this range, small variations in the sinus cycle length result in very different entrainment rhythms (Fig. 3D), as seen in the clinical patient (Fig 2C).

**Fig. 3:**
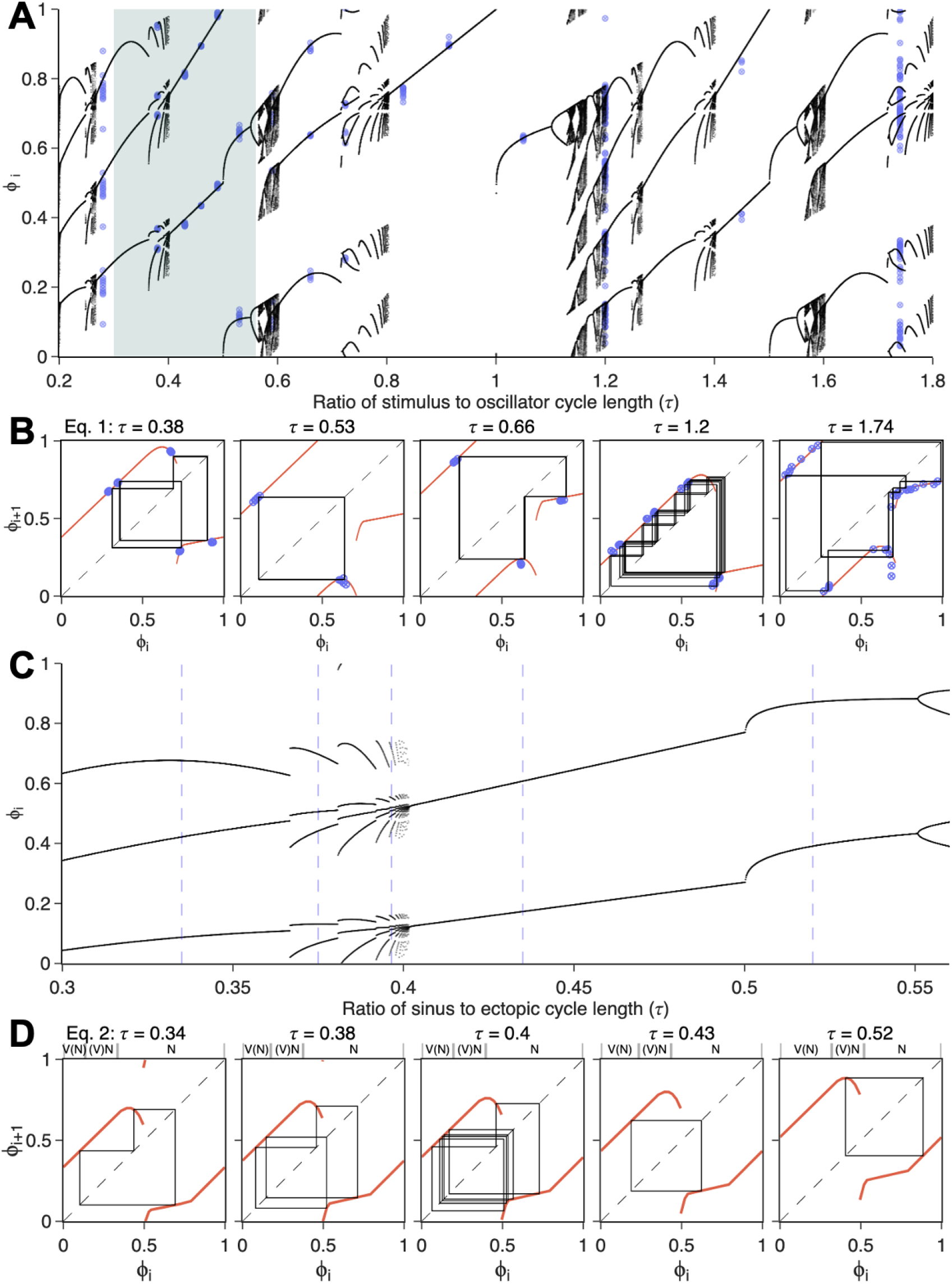
Dynamical regimes observed in experimental and clinical data, explained by bifurcation analysis of mathematical models. (**A**) Bifurcation diagram of the periodically-forced oscillator model (Eq. 1) using the experimentally fitted PRC from Fig. 1(B). Blue dots mark experimental data. The shaded region indicates the range of values observed in the patient data. (**B**) Cobweb plots for Eq.1 at selected values of studied in experiments. The return map is shown in red; model iterations after transients are shown in black; experimental data are overlaid in blue. The identity line, along which, is shown as a dashed line. (**C**) Bifurcation diagram of the modulated parasystole model (Eq. 2), using the same experimentally fitted PRC, plotted over the range of values observed in the patient record. Vertical blue lines indicate specific values corresponding to the clinical segments shown in Fig. 2C. (**D**) Corresponding cobweb plots for Eq. 2 for each value. The three regions above the graph indicate different contexts for the sinus beat depending on its phase. V(N), blocked sinus beat preceded by an ectopic beat; (V)N, sinus beat preceded by a blocked ectopic beat; N, single sinus beat.

These simple reductionist methods highlight universal properties of periodically stimulated oscillators in experiments and people. The arrhythmias generated by an ectopic focus in the heart display a wide range of dynamics that only become obvious when considering the bifurcation analysis based on the PRC. The analysis not only describes the dynamics but also suggests ways to manipulate these rhythms to obtain simple locking regions in which ectopic firing is reduced or abolished. In the following sections, we explore biophysical methods to modify the PRC and how these modifications can be used to reduce ectopic firing across the models.

### Ion current contributors to the PRC in a computational model of hiPSC-CM

We carry out simulations using a computational model of a hiPSC-CM (*32*). The ionic currents modelled are described in Fig. 4A. To construct the PRC, in an AP cycle we deliver a stimulus of 10 ms and 1.5 × 10^−10^A, probing the cycle at 10 ms increments (Fig. 4B). While most stimuli delivered in the first 70% of the cycle have minimal effect, stimuli delivered early in the resting phase lead to a cycle length prolongation. At phases greater than about 0.7, there is an abrupt transition (discontinuity) in response to stimuli; these stimuli trigger an AP. This transition is associated with an all-or-none AP response as a critical voltage is crossed (*27*).

**Fig. 4:**
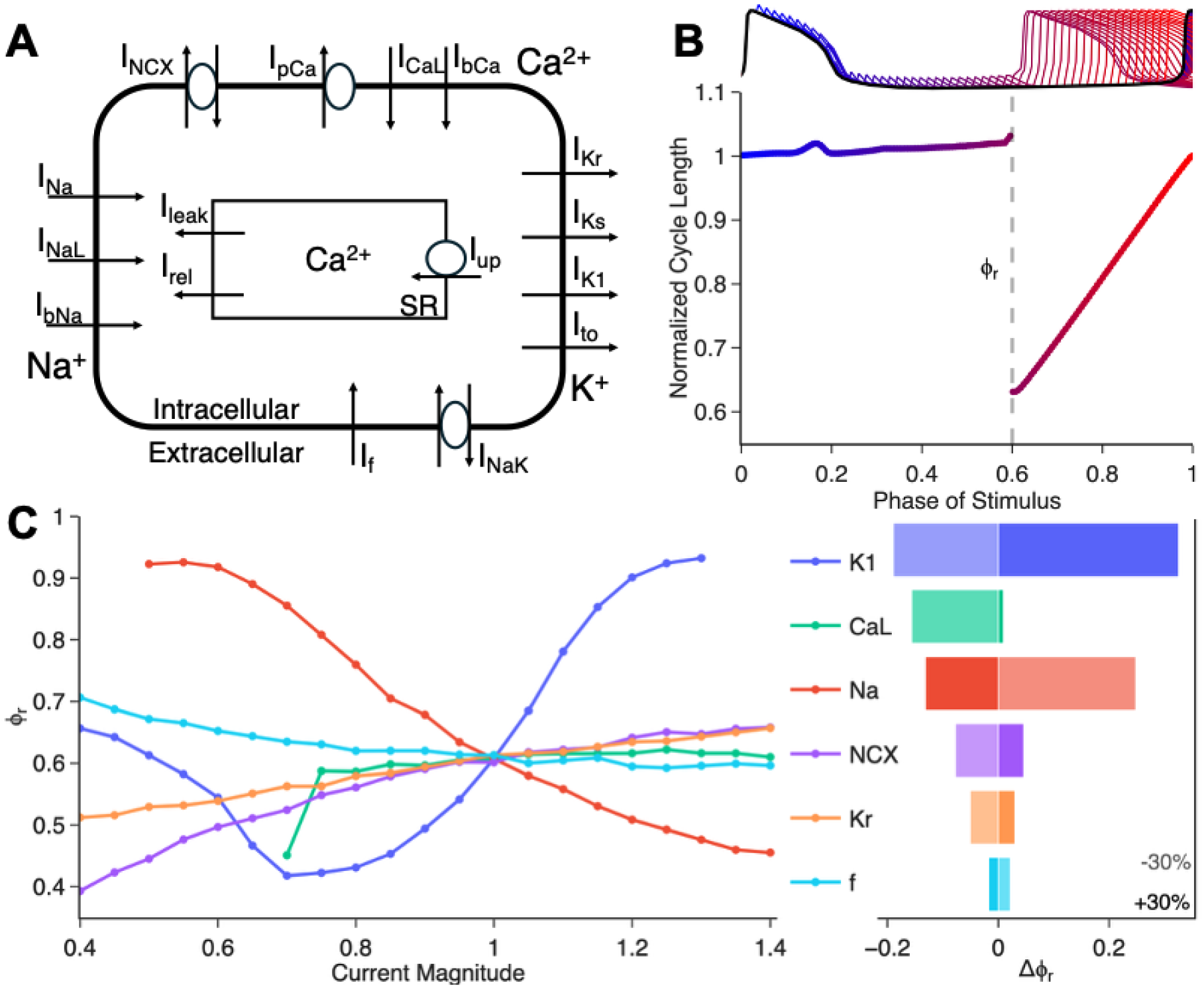
Phase resetting in a computational model for human iPSC-CMs. (**A**) Schematic representation of the ionic currents in the human iPSC-CM model (Paci et al, 2020). (**B**) Top: Trace of a full action potential cycle (black), overlaid with action potentials evoked by stimuli applied at incrementally increasing times (ever 30 ms; traces color-coded by stimulus phase). Bottom: Corresponding phase response curve (PRC). **(C)** Sensitivity of the PRC discontinuity () to ionic current magnitudes (magnitude of 1 corresponds to default value). Left: as a function of current magnitude. Right: Change in following a change in the conductance of each current (shaded: -30%, opaque: +30%).

The location of the discontinuity *ϕ*_*r*_ plays a crucial role in determining the effects of periodic stimulation; decreasing *ϕ*_*r*_ increases the range of stimulus frequencies that lead to simple locking patterns. We examined the role of several ionic currents in the location of this discontinuity, Fig. 4C. In the left panel, each current was varied in strength from 0.4x to 1.4x the default conductance magnitude in the model (1x). The fast sodium current I_Na_ has a strong effect on the PRC, decreasing *ϕ_r_* as the conductance of I_Na_ is increased. The inward rectifier current I_K1_ has a powerful non-monotonic effect on the PRC. An increase from default values can prevent early triggering, while a decrease can significantly facilitate early triggering. Further decrease in I_K1_, however also prevents AP triggering. This is consistent with the complex and context-dependent role of I_K1_ on excitability (*33*). Increases in the L-type calcium current I_CaL_, the Na/Ca exchanger current I_NCX_, the rapid delayed rectifier I_Kr_, and the funny/pacemaking current I_f_ have little effect on *ϕ*_*r*_. However, reducing any of these currents decreases *ϕ*_*r*_, except for I_f_ which has small effect in the opposite direction. In the right panel of Fig. 4C, we show the change in *ϕ*_*r*_ at + or – 30% conductance for each current. Overall, the most significant currents in determining the location of the discontinuity in this model are I_K1_, I_Na_, and I_CaL_. Furthermore, the *in silico* PRC simulations also helped identify environmental factors that decrease *ϕ*_*r*_, helping trigger an early AP (fig. S8).

### Entrainment zones infer strategies to suppress PVCs

Figure 5 illustrates entrainment zones across the different models, and how they can guide strategies to suppress premature ventricular complexes (PVCs). For the periodically-forced oscillator model (Eqn. 1) with a piecewise linear PRC (an approximation to the PRC of the hiPSC-CM computational model, see materials and methods), distinct entrainment zones arise across the *τ* - *ϕ*_*r*_ parameter space (panel A). Notably, smaller values of *ϕ*_*r*_ result in wider low-period locking zones. Similar entrainment zones are present in the hiPSC-CM computational model, where varying sodium channel conductance has a similar effect (panel B). When the oscillator model is extended to include the experimentally derived PRC (panel C) and then incorporated into a modulated parasystole model with patient-derived parameters (panel D), the global structure of the locking zones persists with slight changes based on the nonlinearity introduced in the PRC. The white line indicates the range of *τ* values observed in the patient record. The PVC burden is computed from simulations of the modulated parasystole model over the same parameter range (Panel E). Under the current value of *t*_lag_ fitted to the patient, variations in *ϕ*_*r*_ alone do not provide a reduction in PVC burden. However, when *t*_lag_ is reduced (panel F), decreasing *ϕ*_*r*_ leads to fewer PVCs, as low-period locking becomes favored. In this regime, entrainment of the ectopic focus suppresses PVCs by timing them to fall consistently in the refractory period of the heart. Movement in parameter space toward these low-period zones (indicated by the white arrow) suggests a potential strategy for PVC suppression. This raises the possibility that modifying the PRC, for example through pharmacological interventions, could shift the system into more favorable locking zones and thereby reduce ectopic activity.

**Fig. 5:**
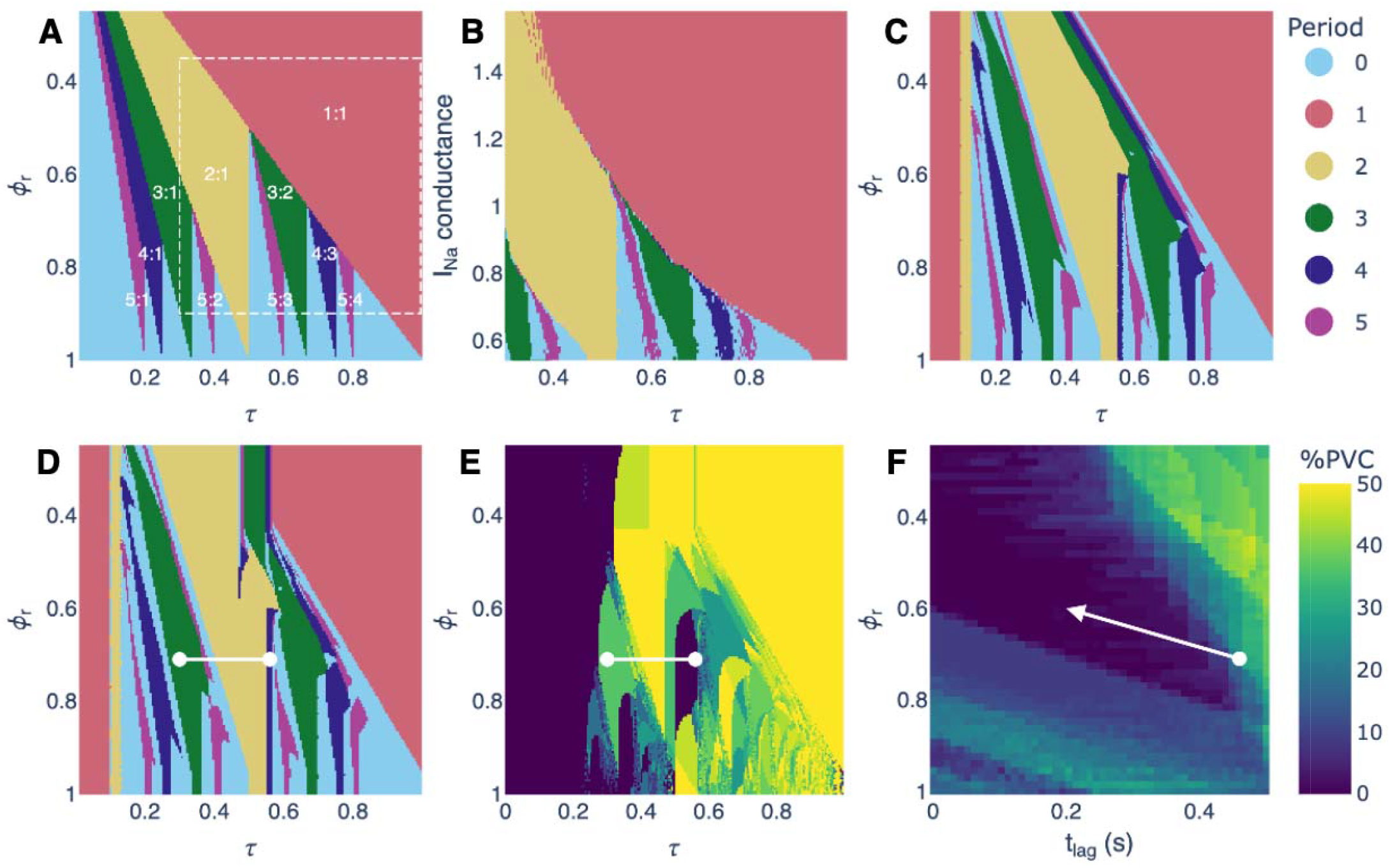
Locking zones across different models and an implied strategy for suppressing PVCs. (**A)** Locking zones in the oscillator model with a piecewise linear PRC, shown as a function of and. Inset numbers indicate the locking ratio N:M, and colors represent the period N; “Period 0” refers to aperiodicity or N. The dashed box highlights the region also examined in the ionic model. (**B**) Locking zones in the ionic model as a function of sodium channel conductance. Similar locking regions appear in the oscillator model with a linear PRC (dashed box). (**C**) Locking zones in the oscillator model using the empirical PRC. (**D**) Locking zones in the modulated parasystole model, using the same PRC and other parameters derived from patient data. The white line indicates the range of values observed in the patient record. (**E**) Percentage of PVCs observed in the modulated parasystole model fit with patient data. (**F**) Percentage of PVCs averaged over, shown as a function of and (conduction time into and out of the ectopic focus). The white dot marks parameter values fit to the patient. The white arrow indicates the direction associated with a reduced PVC burden.

The locking zones in these models illustrate how small changes in parameter values can lead to dramatic changes in dynamics. To reproduce the observed rhythms across a patient record, it was necessary to allow parameters in the model to vary, consistent with the notion that physiological factors such as conduction velocity, autonomic tone, and excitability fluctuate over the day. This non-stationarity may help explain why patients with frequent PVCs sometimes present without PVCs on the day of their scheduled ablation.

## Discussion

We provide experimental characterization of PRCs for hiPSC-CM spheroids, using optogenetic stimulation and optical mapping. These PRCs share morphological similarities with curves from chick embryo heart cells (*34*) and canine Purkinje fibers (*9*), confirming universality of such responses. PRCs from an *in silico* model of these hiPSC-cardiomyocytes reveal the contribution of various ionic currents to phase resetting. There are several important applications of the empirically derived PRCs and the developed parasystole models.

First, as illustrated in our study, the empirical human cardiac PRC (from hiPSC-CM spheroids) captured the dynamics of parasystole in patients with recurrent PVCs. After automatic estimation of a conduction delay (to reflect the distance between the sinus node and the ectopic site from patient ECGs), the empirical PRC modeled the model dynamics in long patient records. Recent work supports the observation that a high frequency of PVCs with parasystolic signature can result from conduction system abnormalities. Further, such parasystole is highly correlated with future occurrence of ventricular fibrillation in cardiomyopathy patients (*35*). While PVCs are generally benign, their frequency is associated with potential induction of lethal arrhythmias. The mathematical model derived here may be useful in estimating the site of origin and severity of the PVCs (Purkinje vs. other) (*36, 37*) for early stratification based on the estimated delay, and in developing strategies for prevention of dangerous arrhythmias.

A second potential area of application of the PRC analysis is the engraftment of hiPSC-CM for heart regeneration. Engrafted hiPSC-CMs behavior is consistent with a stimulated oscillator (*38*). At least eight clinical trials are underway in Japan, Germany and China to deliver hiPSC-CMs or engineered constructs of such cells in patients with heart failure or other cardiac pathology (*12, 14*). Engraftment of these cells in large animal models often was associated with transient engraftment arrhythmias within the first two weeks while the graft is loosely coupled to the donor heart. Large-scale tissue simulations focused exclusively on the graft-host electrical coupling affecting the engraftment arrhythmia propensity (*18, 19, 39*). As such coupling strengthens over time, the engraftment arrhythmias subside. In addition, suppression of the ectopic activity from the graft reduced the incidence of lethal arrhythmias originating from this ectopic focus (*40*).

There is a desire to increase the safety of hiPSC-CM transplantation by engineering the graft to have anti-arrhythmic properties upon transplantation or by administering pharmacological therapy within the vulnerable window. Our PRC analysis and parasystole modeling offer insights on ion channel targets and can inform ongoing pharmacological treatments (*41*) and gene-editing approaches (*17*) to prevent engraftment arrhythmias. This can be done by revealing critical molecular targets to enlarge the simple locking entrainment zones (Fig. 4-5), by in depth exploration of the dynamics of the system (Fig. 3), and by data-driven personalization for a particular patient (Fig. 2) to devise rhythm control strategies. While large-scale 2D and 3D cardiac tissue modeling with patient-specific geometries is making strides in devising “digital twins” in clinical cardiology (*42*), longer-term dynamics simulations of rhythm disturbances require a different, computationally efficient approach, as showcased here, to fully leverage multi-day patient records for personalized diagnosis and treatment.

## Funding

Canadian Institutes of Health Research and Heart & Stroke Foundation of Canada AB2-190680 (KD)

Fonds de Recherche du Québec—Nature et Technologies postdoctoral fellowship (TMB)

Canadian Institutes of Health Research PJT-169008 (AS)

Government of Canada’s New Frontiers in Research Fund (NFRF) NFRFT-2022-00447 (GB)

Natural Research Sciences and Engineering Research Council of Canada (NSERC) RGPIN-2024-04518 (GB)

Human Frontiers Science Program RGP010/2024 (MEP, EE)

## Author contributions

Conceptualization: KD, TMB, AS, LG, GB, EE

Methodology: KD, TMB, MEP, MWD, ZL, AS, LG, GB, EE

Investigation: KD, TMB, MEP

Visualization: KD, TMB, MEP

Funding acquisition: KD, TMB, AS, GB, EE

Project administration: AS, LG, GB, EE Supervision: AS, LG, GB, EE

Writing – original draft: KD, TMB, MEP, LG, GB, EE

Writing – review & editing: KD, TMB, MEP, MWD, ZL, AS, LG, GB, EE

## Data and materials availability

All data, code, and materials used in the analysis are deposited in the GitHub repository github.com/KhadyDiagne/PRC-of-hiPSC-CM.

## Supplementary Materials

Materials and Methods Figs. S1 to S8

Tables S1 to S3

## References and Notes

1. B. Z. Jia, Y. Qi, J. D. Wong-Campos, S. G. Megason, A. E. Cohen, A bioelectrical phase transition patterns the first vertebrate heartbeats. Nature 622, 149–155 (2023).

2. A. T. Winfree, Sudden cardia death: a problem in topology. Sci.Am. 248, 144–147, 160 (1983).

3. M. C. Mackey, L. Glass, Oscillation and chaos in physiological control systems. Science 197, 287–289 (1977).

4. M. R. Guevara, L. Glass, A. Shrier, Phase locking, period-doubling bifurcations, and irregular dynamics in periodically stimulated cardiac cells. Science 214, 1350–1353 (1981).

5. T. R. Chay, Y. S. Lee, Phase resetting and bifurcation in the ventricular myocardium. Biophys J 47, 641–651 (1985).

6. J. Jalife, C. Antzelevitch, G. K. Moe, The case for modulated parasystole. Pacing Clin.Electrophysiol. 5, 911–926 (1982).

7. A. T. Winfree, Phase control of neural pacemakers. Science 197, 761–763 (1977).

8. G. K. Moe, J. Jalife, W. J. Mueller, B. Moe, A mathematical model of parasystole and its application to clinical arrhythmias. Circulation 56, 968–979 (1977).

9. J. Jalife, G. K. Moe, A biologic model of parasystole. Am.J.Cardiol. 43, 761–772 (1979).

10. K. Diagne et al., Rhythms from two competing periodic sources embedded in an excitable medium. Phys Rev Lett 130, 028401 (2023).

11. M. Courtemanche, L. Glass, M. D. Rosengarten, A. L. Goldberger, Beyond pure parasystole: promises and problems in modeling complex arrhythmias. Am J Physiol 257, H693–H706 (1989).

12. T. Sugiura, D. C. Shahannaz, B. E. Ferrell, Current status of cardiac regenerative therapy using induced pluripotent stem cells. Int J Mol Sci 25, (2024).

13. A. F. Jebran et al., Engineered heart muscle allografts for heart repair in primates and humans. Nature 639, 503–511 (2025).

14. W. Yan et al., Stem cell-based therapy in cardiac repair after myocardial infarction: Promise, challenges, and future directions. Journal of Molecular and Cellular Cardiology 188, 1–14 (2024).

15. Y. Shiba et al., Allogeneic transplantation of iPS cell-derived cardiomyocytes regenerates primate hearts. Nature 538, 388–391 (2016).

16. R. Romagnuolo et al., Human embryonic stem cell-derived cardiomyocytes regenerate the infarcted pig heart but induce ventricular tachyarrhythmias. Stem cell reports 12, 967–981 (2019).

17. S. Marchiano et al., Gene editing to prevent ventricular arrhythmias associated with cardiomyocyte cell therapy. Cell Stem Cell 30, 741 (2023).

18. C. E. Gibbs, P. M. Boyle, Population-based computational simulations elucidate mechanisms of focal arrhythmia following stem cell injection. J Mol Cell Cardiol 204, 5–16 (2025).

19. C. E. Gibbs et al., Graft-host coupling changes can lead to engraftment arrhythmia: a computational study. J Physiol 601, 2733–2749 (2023).

20. C. Antzelevitch, M. J. Bernstein, H. N. Feldman, G. K. Moe, Parasystole, reentry, and tachycardia: a canine preparation of cardiac arrhythmias occurring across inexcitable segments of tissue. Circulation 68, 1101–1115 (1983).

21. Z. Jia et al., Stimulating cardiac muscle by light: cardiac optogenetics by cell delivery. Circ Arrhythm Electrophysiol 4, 753–760 (2011).

22. A. Klimas, G. Ortiz, S. C. Boggess, E. W. Miller, E. Entcheva, Multimodal on-axis platform for all-optical electrophysiology with near-infrared probes in human stem-cell-derived cardiomyocytes. Prog Biophys Mol Biol 154, 62–70 (2020).

23. E. Entcheva, M. W. Kay, Cardiac optogenetics: a decade of enlightenment. Nature reviews. Cardiology 18, 349–367 (2021).

24. Y. L. Huang, A. S. Walker, E. W. Miller, A photostable silicon rhodamine platform for optical voltage sensing. J Am Chem Soc 137, 10767–10776 (2015).

25. D. H. Perkel, J. H. Schulman, T. H. Bullock, G. P. Moore, J. P. Segundo, Pacemaker neurons: effects of regularly spaced synaptic input. Science 145, 61–63 (1964).

26. L. Glass, M. R. Guevara, J. Belair, A. Shrier, Global bifurcations of a periodically forced biological oscillator. Phys Rev A 29, 1348–1357 (1984).

27. T. Krogh-Madsen, L. Glass, E. J. Doedel, M. R. Guevara, Apparent discontinuities in the phase-resetting response of cardiac pacemakers. J Theor Biol 230, 499–519 (2004).

28. V. C. Kowtha, A. Kunysz, J. R. Clay, L. Glass, A. Shrier, Ionic mechanisms and nonlinear dynamics of embryonic chick heart cell aggregates. Prog Biophys Mol Biol 61, 255–281 (1994).

29. G. M. Marcus, Evaluation and management of premature ventricular complexes. Circulation 141, 1404–1418 (2020).

30. T. M. Bury et al., The inverse problem for cardiac arrhythmias. Chaos 33, (2023).

31. K. Takayanagi et al., Ectopic cycle length estimation from the quantified distribution patterns of ventricular bigeminy and trigeminy. Heart Rhythm O2 2, 138–148 (2021).

32. M. Paci et al., All-optical electrophysiology refines populations of in silico human ipsc-cms for drug evaluation. Biophys J 118, 2596–2611 (2020).

33. J. S. Langen, P. M. Boyle, D. Malan, P. Sasse, Optogenetic quantification of cardiac excitability and electrical coupling in intact hearts to explain cardiac arrhythmia initiation. Sci Adv 11, eadt4103 (2025).

34. J. R. Clay, R. M. Brochu, A. Shrier, Phase resetting of embryonic chick atrial heart cell aggregates. Experiment and theory. Biophys J 58, 609–621 (1990).

35. H. Do Duc et al., Ventricular parasystole in cardiomyopathy patients. JACC: Clinical Electrophysiology 9, 936–948 (2023).

36. W. Escande et al., Malignant Purkinje ectopy induced by sodium channel blockers. Heart Rhythm 19, 1595–1603 (2022).

37. M. Haïssaguerre et al., Idiopathic ventricular fibrillation: role of purkinje system and microstructural myocardial abnormalities. JACC: Clinical Electrophysiology 6, 591–608 (2020).

38. E. Huethorst et al., Evidence for intermittent coupling of intramyocardial small, engineered heart tissues acutely implanted into rabbit myocardium. Cardiovasc Res, (2025).

39. J. K. Yu, J. A. Liang, S. H. Weinberg, N. A. Trayanova, Computational modeling of aberrant electrical activity following remuscularization with intramyocardially injected pluripotent stem cell-derived cardiomyocytes. Journal of Molecular and Cellular Cardiology 162, 97–109 (2022).

40. J. S. Yang, A. R. Ochs, C. E. Gibbs, P. M. Boyle, Computational simulations show proof-of-concept for optogenetic suppression of ectopic activity in cardiac stem cell therapy. Cardiovascular engineering and technology, (2025).

41. K. Nakamura et al., Pharmacologic therapy for engraftment arrhythmia induced by transplantation of human cardiomyocytes. Stem cell reports 16, 2473–2487 (2021).

42. R. Laubenbacher, B. Mehrad, I. Shmulevich, N. Trayanova, Digital twins in medicine. Nat Comput Sci 4, 184–191 (2024).

