## Supplemental Information for "Phase resetting in human stem cell derived cardiomyocytes explains complex cardiac arrhythmias"


**This PDF file includes:**

– Materials and Methods

– Figs. S1 to S8

– Tables S1 to S3

– References

### 1 Materials and Methods

#### 1.1 Phase-resetting curves from human iPSC-CMs

To collect experimental PRCs from human iPSC-CMs we used a two-spheroid model, where an optogenetically-responsive cell spheroid was coupled to a spheroid of unmodified human iPSC-CMs and light pulses were used to perturb the system, while the generated responses (action potentials) were measured optically.

##### 1.1.1 Cell spheroid assembly and pairing

We previously developed an immortalized 293T-HEK cell line expressing ChR2 tagged with yellow fluorescent protein (YFP) (*1*) using the Addgene construct pcDNA3.1/hChR2(H134R)-eYFP (from Karl Deisseroth). ChR2-HEK cells were cultured in Dulbecco's Modified Eagle Medium (DMEM, ATCC) with 10% fetal bovine serum (FBS), 1% 200mM L-Glutamine, and 1% penicillin-streptomycin. After trypsinization, using 0.05% trypsin-disodium ethylenediaminetetraacetic acid (EDTA), spheroids were formed by plating the cell suspension at $\sim 2.5$ to $10 \times 10^3$ cells per well in ultra-low-attachment, round bottom Corning 96-well (#CLS4920, Millipore Sigma) or 384-well (#CLS3830, Millipore Sigma) plates. Spheroids were typically used the following day.

Human induced pluripotent stem-cell-derived cardiomyocytes (hiPSC-CMs) were thawed following the manufacturer's protocol (iCell Cardiomyocytes, #R1017, Fujifilm-Cellular Dynamics). The cells were suspended in plating media and seeded at about 5 to $10 \times 10^3$ cells/well in ultra-low attachment, round bottom plates. 50% of iCell maintenance medium was replaced every 48 hours to maintain spheroid integrity.

One ChR2-HEK spheroid and one hiPSC-CM spheroid were carefully transferred to the same well in a separate Corning 384-well spheroid microplate, avoiding shear forces, and the paired spheroids were incubated for 3 to 24 h to form electrical contact.

##### 1.1.2 Functional experiments with all-optical electrophysiology

Measurements were performed on an inverted Nikon Eclipse Ti2 microscope at 20× magnification. The hiPSC-CM spheroids were labeled with the small-molecule, voltage-sensitive BeRST1 dye at a

2 $\mu$M concentration (generously provided by Evan W. Miller (*2*)), following procedures established in previous studies (*3*). Optogenetic stimulation was done through a digital micromirror device (DMD) (Polygon 4000, Mightex, Toronto, ON, Canada) using 470 nm blue light pulses at 10 ms duration, < 0.15 mW/mm$^2$, at various frequencies to drive the hiPSC-CM spheroids while the illuminated region was covering the ChR2-HEK spheroid. For entrainment experiments, the same setup was used, but pulse durations were adjusted to 5, 10, or 20 ms. Optical voltage imaging with BeRST1 dye involved 660 nm excitation, with emitted light captured with a long pass filter at 700 nm using an iXon Ultra 897 EMCCD camera (Andor Technology Ltd., Belfast, United Kingdom), at 206 fps.

##### 1.1.3 Experimental protocols

To collect data for construction of the PRC, for each spheroid pair, a 40 sec recording of spontaneous activity was taken initially to calculate the hiPSC-CM spheroid's spontaneous rate. Using this spontaneous rate, the stimulus frequency was defined by solving

$$\mathrm{R} = \frac{\text{Spontaneous frequency (Hz)}}{\text{Stimulus Frequency (Hz)}} \tag{1}$$

where $R$ is a user-defined ratio. A subsequent recording was then taken consisting of three phases: the first 40 sec captured spontaneous activity (no excitation), the next 10 min applied optical excitation to the stimulation region at the calculated frequency, and the final 40 sec recorded spontaneous activity to give the excitable cells time to recover between each tested frequency. Voltage activity was recorded from the sensing region of the hiPSC-CM spheroid for the entire duration. This process, including recalculating the spontaneous rate, was conducted for each value of $R$. After all tests, one final spontaneous recording was taken. These recordings were performed for $n$ = 6 spheroid pairs with all optical pulses set to 10 ms durations.

To collect entrainment data, a similar protocol was used with some differences. Based on an initial 40 sec record of spontaneous frequency, 13-17 various pacing frequencies were applied - below, above, or close to the spontaneous rate. Each recording consisted of three phases: 30 sec of spontaneous activity, 1 min of optical excitation, then a final 30 sec of spontaneous activity for recovery between frequencies. Additional 40 sec spontaneous recordings were performed periodically to check the spontaneous rate and a final spontaneous recording was taken after all measurements were completed. These recordings were conducted on $n = 1$ spheroid pair with data collected at optical pulse durations of 5, 10, and 20 ms. Phase resetting data was also obtained for this spheroid pair at all three pulse durations using the methods described previously, with 5 min of optical stimulation recorded instead of 10 min.

###### 1.1.4 Fitting a polynomial equation to experimental PRCs

We studied entrainment and phase-resetting in six spheroid pairs. Their PRCs are shown in Fig. S1. We model the PRCs using the function

$$g(\phi) = \begin{cases} 1 & 0 \le \phi < \phi_1, \\ 1 + A(\phi - \phi_1)^4 & \phi_1 \le \phi < \phi_r, \\ B(\phi - \phi_2)^2 + C & \phi_r \le \phi < \phi_3, \\ 1 + S(\phi - 1) & \phi_3 \le \phi < 1, \end{cases} \tag{2}$$

which is a piecewise-defined polynomial equation designed to be as simple as possible while capturing the main features observed in the experimental PRCs. The parameters $A$, $\phi_r$, $B$ and $S$ are fit to the experimental data (see Table S1) using a nonlinear least-squares optimization algorithm (Trust Region Reflective method). Specifically, $A$ controls the magnitude of the delay branch, $\phi_r$ defines the point of discontinuity between delay and advance, $B$ controls the curvature of the hook in the initial portion of the advance branch, and $S$ is the slope of the later portion of the advance branch. The parameters $\phi_1$, $\phi_2$, $\phi_3$ and $C$ are auxiliary: they are not fit directly but impose structural constraints and ensure continuity and smoothness. We set $\phi_1 = \phi_r - 0.25$ and $\phi_2 = \phi_r + 0.04$. The parameters $\phi_3$ and $C$ are uniquely determined to ensure continuity of both the function and its derivative at $\phi = \phi_3$, given by

$$\phi_3 = \phi_2 + S/(2B), \tag{3}$$

$$C = 1 - S(1 - \phi_3) - B(\phi_3 - \phi_2)^2. \tag{4}$$

The PRC shown in Fig. S1A was used for the analysis in the main text. All six datasets showed qualitatively similar features: no resetting at early phases, a delay at intermediate phases, and apparent discontinuity leading to an advance at later phases. A hook in the early portion of the advancing branch was also visible in each dataset. The most prominent variation among the spheroids was the point of discontinuity $\phi_r$. The hiPSC-CM spheroids in panels A–C have later points of discontinuity (0.71, 0.57, and 0.57, respectively) compared to those in panels D–F (0.42, 0.40, and 0.38, respectively). This suggests that the latter group is more readily entrained, possibly due to stronger effective stimuli. The magnitude of the stimulus influences the PRC (*4*). It is likely that variability in the contact area between the ChR2-HEK and hiPSC-CM spheroids, as well as differences in spheroid size, contributed to these variations even under constant pulse duration.

#### 1.2 Parasystole and resetting in clinical ECGs

##### 1.2.1 Selection criteria and preprocessing of Holter ECG recordings from patients

We have multi-day Holter recordings from 53 patients enrolled in the British Columbia PVC Registry, all of whom exhibit $> 5\%$ premature ventricular complexes (PVCs). Patients were included for further analysis according to the criteria shown in Fig. S2. For consistency with our modeling assumptions, we included only those patients whose PVCs were mostly of a single morphology, consistent with PVCs originating from a single ectopic focus. Additionally, we flagged patients as potential candidates for parasystole by applying a mask to identify coupling intervals that systematically "march in" or "march out."

**Mask to identify cycling coupling intervals.** To identify dynamics consistent with parasystole, we designed a rule-based mask to detect sequences in which the coupling interval progressively lengthens ("marching in") or shortens ("marching out"). A sequence was classified as *marching in* if it satisfied all of the following criteria:

1. alternating ectopic and sinus beats;
2. coupling intervals that progressively lengthen;
3. compensatory pauses that progressively shorten;
4. two consecutive sinus beats that terminate the sequence of alternating ectopic and sinus beats.

Similarly, a sequence was classified as *marching out* if it satisfied:

1. alternating ectopic and sinus beats;
2. coupling intervals that progressively shorten;

3. compensatory pauses that progressively lengthen;
4. two consecutive sinus beats prior to the sequence of alternating ectopic and sinus beats.

These masks were applied to each patient record to isolate segments exhibiting cycling coupling intervals suggestive of parasystole. Representative examples of both "marching in" and "marching out" patterns are shown in Fig. S3. A patient was selected for further analysis if their record contained at least three consecutive sequences of marching in or marching out. Based on this criterion, 7 patients (13% of the cohort) were included for further analysis.

##### 1.2.2 Mathematical model for modulated parasystole

**An early model for modulated parasystole.** We build on an early model for modulated parasystole by Courtemanche et al. (*5*). This model assumes a sinus pacemaker with period $t_s$, an ectopic pacemaker with period $t_e$ a refractory period $\theta$, and a phase response curve for the ectopic pacemaker $g(\phi)$. The ectopic beat is blocked if it lands within the refractory period of the sinus beat. The sinus beat following an ectopic beat is always blocked. The model is formulated as follows. Let $\phi_i$ be the phase of the sinus beat within the ectopic cycle. Then one can determine the context in which it occurred, as outlined below:

| Phase Range | Beat Sequence | Description |
|---|---|---|
| $0 \leq \phi < \frac{t_s-\theta}{t_e}$ | V(N) | The sinus beat is blocked due to a preceding ectopic beat. |
| $\frac{t_s-\theta}{t_e} < \phi < \frac{t_s}{t_e}$ | (V)N | The sinus beat is expressed and preceded by a blocked ectopic beat. |
| $\frac{t_s}{t_e} \leq \phi < 1$ | N | The sinus beat is expressed. |

If a sinus beat is blocked, there is no modulation of the ectopic pacemaker. The next phase is then obtained by adding the normalized sinus period $t_s/t_e$. If a sinus beat is expressed, there is modulation of the ectopic pacemaker, resulting in a shift of the phase by $1 - g(\phi)$. This yields the following difference equation for the phase

$$\phi_{i+1} = \begin{cases} \phi_i + \frac{t_s}{t_e} \pmod 1 & 0 \le \phi_i < \frac{t_s - \theta}{t_e}, \\ \phi_i + \frac{t_s}{t_e} + 1 - g(\phi) \pmod 1 & \frac{t_s-\theta}{t_e} \le \phi_i < 1, \end{cases} \tag{5}$$

where $0 \le \phi_i < 1$.

**Extension to include a conduction time into and out of the ectopic focus.** Electrophysiological studies have reported a conduction delay between the initial firing of the ectopic focus and the subsequent PVC (*6*). We denote this delay as $t_{\text{out}}$. Similarly, we assume a conduction delay between a sinus beat and the resetting of the ectopic pacemaker, which we denote $t_{\text{in}}$ (Fig. S4). Assumption of these conduction delays requires modification to the model equations. Let $\phi$ be the phase of the sinus beat in the ectopic cycle as measured by the timing of the ectopic and sinus beat on an ECG (Fig. S4). Now, the elapsed time between firing of the focus and resetting of the focus is

$$t' = t_{\text{out}} + \phi t_e + t_{\text{in}} \tag{6}$$

which corresponds to a phase in the cycle of the ectopic pacemaker of

$$\phi' = \frac{t'}{t_e} = \phi + \frac{t_{\text{lag}}}{t_e} \tag{7}$$

where $t_{\text{lag}} = t_{\text{in}} + t_{\text{out}}$. To incorporate this property into the model, we modify the argument of the phase response curve accordingly. Other aspects remain unchanged. The model with conduction delay is given by

$$\phi_{i+1} = \begin{cases} \phi_i + \frac{t_s}{t_e} \pmod 1 & 0 \le \phi_i < \frac{t_s - \theta}{t_e}, \\ \phi_i + \frac{t_s}{t_e} + 1 - g\left(\phi + \frac{t_{\text{lag}}}{t_e}\right) \pmod 1 & \frac{t_s-\theta}{t_e} \le \phi_i < 1, \end{cases} \tag{8}$$

where $0 \le \phi_i < 1$.

**Model variant that permits interpolated beats.** Previous models (*5, 7*) assume that the sinus beat immediately following an ectopic beat is always blocked. This is not always the case in clinical data, where a patient can experience an interpolated PVC—a PVC that does not lead to a block of the subsequent sinus beat. In this model variant, we assume that the sinus beat is only blocked if it occurs during a refractory period following the ectopic beat. We take this refractory period to be $\theta$, the same value that is assigned to the refractory period of the sinus beats.

Note that if $t_s < 2\theta$, a sinus beat following an ectopic necessarily lands in the refractory period. Therefore, for this range of sinus rates, the model is identical to Eqn. (8). However, for slower sinus rates $t_s > 2\theta$, there are four different outcomes depending on the phase, outlined below:

| Phase Range | Beat Sequence | Description |
|---|---|---|
| $0 \le \phi < \frac{\theta}{t_e}$ | V(N) | An ectopic beat followed by a blocked sinus beat. |
| $\frac{\theta}{t_e} \le \phi < \frac{t_s-\theta}{t_e}$ | VN | An (interpolated) ectopic beat followed by a sinus beat. |
| $\frac{t_s-\theta}{t_e} \le \phi < \frac{t_s}{t_e}$ | (V)N | A blocked ectopic beat followed by a sinus beat. |
| $\frac{t_s}{t_e} \le \phi < 1$ | N | An additional sinus beat in the ectopic cycle |

In this model, phase resetting only occurs for $\phi \ge \min(\theta/t_e, (t_s-\theta)/t_e)$. The difference equation model that permits interpolated beats is

$$\phi_{i+1} = \begin{cases} \phi_i + \frac{t_s}{t_e} \pmod 1 & 0 \le \phi_i < \min\left(\frac{\theta}{t_e}, \frac{t_s-\theta}{t_e}\right), \\ \phi_i + \frac{t_s}{t_e} + 1 - g\left(\phi + \frac{t_{\text{lag}}}{t_e}\right) \pmod 1 & \min\left(\frac{\theta}{t_e}, \frac{t_s-\theta}{t_e}\right) \le \phi_i < 1, \end{cases} \tag{9}$$

where $0 \le \phi_i < 1$.

##### 1.2.3 Construction of PRC from clinical ECG

We use segments of bigeminy (a single sinus beat between two ectopic beats) and trigeminy (two sinus beats between two ectopic beats) to construct an approximation of the PRC from clinical data (*8*). Notation for beat-to-beat intervals is shown in Fig. S5. We make the following assumptions: (i) the intervening sinus beats occur during a single cycle of the ectopic pacemaker; (ii) during trigeminy, the second sinus beat shortens the ectopic cycle. We let $g$ denote the PRC that maps beat intervals according to

$$\text{VV}_1 = g(\text{VN}_1). \tag{10}$$

**Construction from bigeminy data.** During bigeminy, we collect data in the form of $(\text{VN}_1, \text{VV}_1)$, which enables us to approximate $g$ over a limited range of $\text{VN}_1$ values. In clinical records, this relationship is typically a positive linear trend. Assuming a linear form,

$$\mathrm{VV}_1 = S * \mathrm{VN}_1 + C, \tag{11}$$

we estimate the slope $S$ and intercept $C$ using linear regression. In addition, we assume that when $\mathrm{VN}_1 = \mathrm{VV}_0 - t_{\mathrm{lag}}$, the sinus stimulus lands at the end of the ectopic cycle (recall Fig. S4), resulting in no resetting ($\mathrm{VV}_1 = \mathrm{VV}_0$). Under this condition

$$\mathrm{VV}_0 = g(\mathrm{VV}_0 - t_{\mathrm{lag}}) \implies C = (1 - S)\mathrm{VV}_0 + S\, t_{\mathrm{lag}}, \tag{12}$$

which provides a relationship between $\mathrm{VV}_0$ and $t_{\mathrm{lag}}$ determined from data.

**Construction from trigeminy data.** During trigeminy, we collect data in the form of $(\mathrm{VN}_1, \mathrm{VN}_2, \mathrm{VV}_2)$. We would like to infer $\mathrm{VV}_1$ from these data in order to uncover a wider range of the PRC. The first sinus beat modifies the ectopic cycle length by $\Delta T = \mathrm{VV}_1 - \mathrm{VV}_0$. Resetting from the second sinus beat then satisfies

$$\mathrm{VV}_2 = g(\mathrm{VN}_2 - \Delta T) + \Delta T. \tag{13}$$

Assuming the second sinus beat shortens the ectopic cycle length according to Eqn. 11, we have

$$\mathrm{VV}_2 = S(\mathrm{VN}_2 - \Delta T) + C + \Delta T. \tag{14}$$

Substituting the expression for $C$ from Eqn. 12 and rearranging yields

$$\mathrm{VV}_1 = \frac{\mathrm{VV}_2 - S(\mathrm{VN}_2 + t_{\mathrm{lag}})}{1 - S}. \tag{15}$$

Since we have estimates for $S$ and $t_{\mathrm{lag}}$ (see below), and $\mathrm{VV}_2$ and $\mathrm{VN}_2$ are measured directly from trigeminy sequences, we can infer $\mathrm{VV}_1$ and augment the dataset used to construct the PRC.

**Phase response curve of the ectopic pacemaker.** Above, we have written the phase response curve in terms of beat-to-beat intervals on the ECG. The phase response curve of the ectopic pacemaker can be obtained by rescaling to get the phase as follows:

$$\phi = \frac{\mathrm{VN}_1 + t_{\mathrm{lag}}}{t_e}. \tag{16}$$

##### 1.2.4 Parameter estimation for the modulated parasystole model

The parameters of the modulated parasystole model are listed in Table S2, along with the methods used to estimate them from the clinical and experimental datasets. The sinus cycle length ($t_s$) is estimated as the mean interbeat interval in a 30-second section of a recording. The ectopic cycle length ($t_e$), the lag time ($t_{\text{lag}}$) and PRC discontinuity ($\phi_r$) are determined by fitting the model to a 30-second segment of data identified as exhibiting a cycling coupling interval (Fig. S3). These parameters are chosen to minimize the mean absolute error between the clinical and model-generated interbeat intervals. The PRC resetting slope ($S$) is determined by the slope in the plot of VV intervals against VN intervals (Fig. S6). The refractory period ($\theta$) is determined from the lower bound of NV intervals at each heart rate (Fig. S7). The remaining parameters ($m$, $p$, $r$, $N$), which govern the shape of the PRC, are obtained from experimental data. Individual parameter values for each patient are reported in Table S3.

Using the fitted parameter values, we evaluate the model's ability to reproduce other 30-second segments from the same recording. The analysis is restricted to segments that do not contain unidentified beats. In this evaluation, the model is not simulated with a fixed sinus cycle length ($t_s$), but instead with a sequence of sinus cycle lengths obtained from averaging the patient interbeat intervals over a rolling window of two beats. To assess goodness of fit, we use the two-tailed Kolmogorov-Smirnov test, which evaluates whether two distributions are significantly different. The model is considered a good fit if the Kolmogorov–Smirnov test does not detect a statistically significant difference between the interbeat interval distributions of the patient and the model ($p > 0.05$). The proportion of 30-second segments that meet this criterion is reported in Table S3.

#### 1.3 Entrainment in an ionic model of a human iPSC-CM

We investigated the phase resetting curves (PRCs) from an ionic computational model of human cardiomyocyte by Paci et al., that was constrained by multiple experimental voltage and calcium records obtained by all-optical electrophysiology from the same ventricular-like hiPSC-CMs as used here (*9*). This model follows the classic Hodgkin-Huxley formulation and includes the cytosol and sarcoplasmic reticulum compartments. The MATLAB implementation is made available by the authors. We simulate the model at 37°C with external ionic concentrations of $[Na^+]_o$ = 151 mM,

$[K^+]_o$ = 5.4 mM, and $[Ca^{2+}]_o$ = 1.8 mM, and internal initial ionic concentrations of $[Na^+]_i$ = 9.2 mM, $[K^+]_i$ = 150 mM, $[Ca^{2+}]_i$ = 0.0002 mM, and $[Ca^{2+}]_{SR}$ = 0.32 mM. Keeping the default parameters of the model, we obtain a cycle length of 1.71 s for the simulated cell.

Using this in silico model, we corroborate that the phase discontinuity shifts to the left and the entrainment is easier for lower temperature, higher stimulus strength and longer stimulus duration, (Fig. S8). Note that for weak or short stimuli, there is no apparent discontinuity in the PRC.

For the locking zones in Figure 5A of the manuscript, we use a piecewise linear equation to model the PRC.

$$g(\phi) = \begin{cases} 1 + a\phi & 0 \leq \phi < \phi_r, \\ 1 + b(\phi - 1) & \phi_r \leq \phi < 1, \end{cases} \tag{17}$$

With parameters $a = 0$, and $b = 1$, this serves as a simple approximation to the PRC obtained in the ionic model.

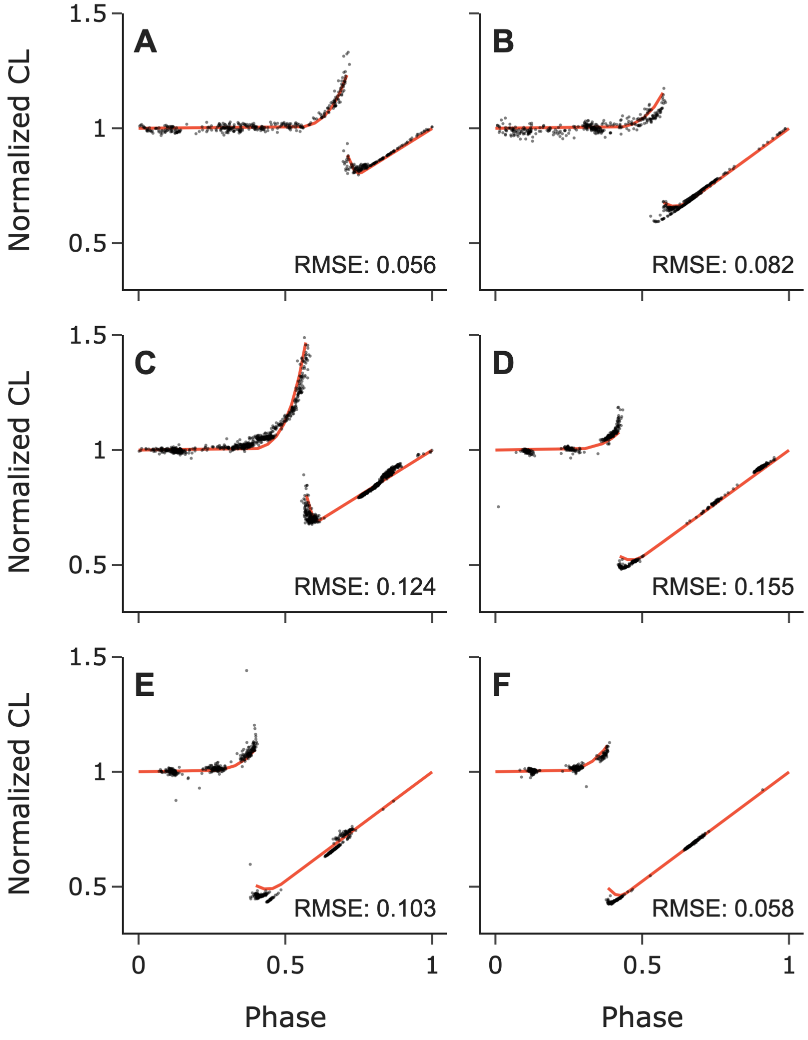


**Figure S1**: **Phase-resetting curves for individual hiPSC-CM spheroids.** The red line shows the fitted PRC function given by Eqn. (2). Optimal parameters were obtained using a nonlinear least-squares algorithm (Trust Region Reflective method). Best-fit parameter values for each aggregate are listed in Table S1. The inset shows the root mean squared error (RMSE) of each fit.

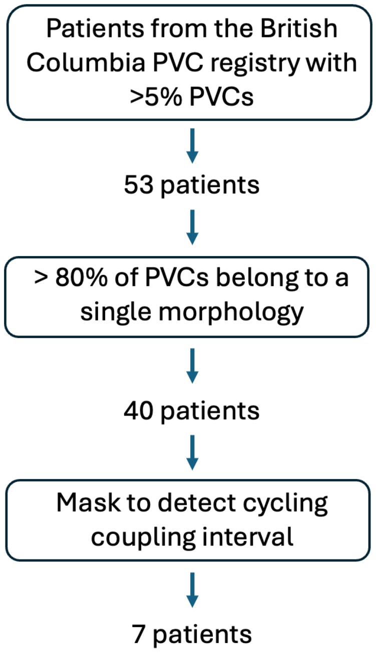


**Figure S2**: Filtering of clinical data based on a the presence of a dominant PVC morphology and a cycling coupling interval.

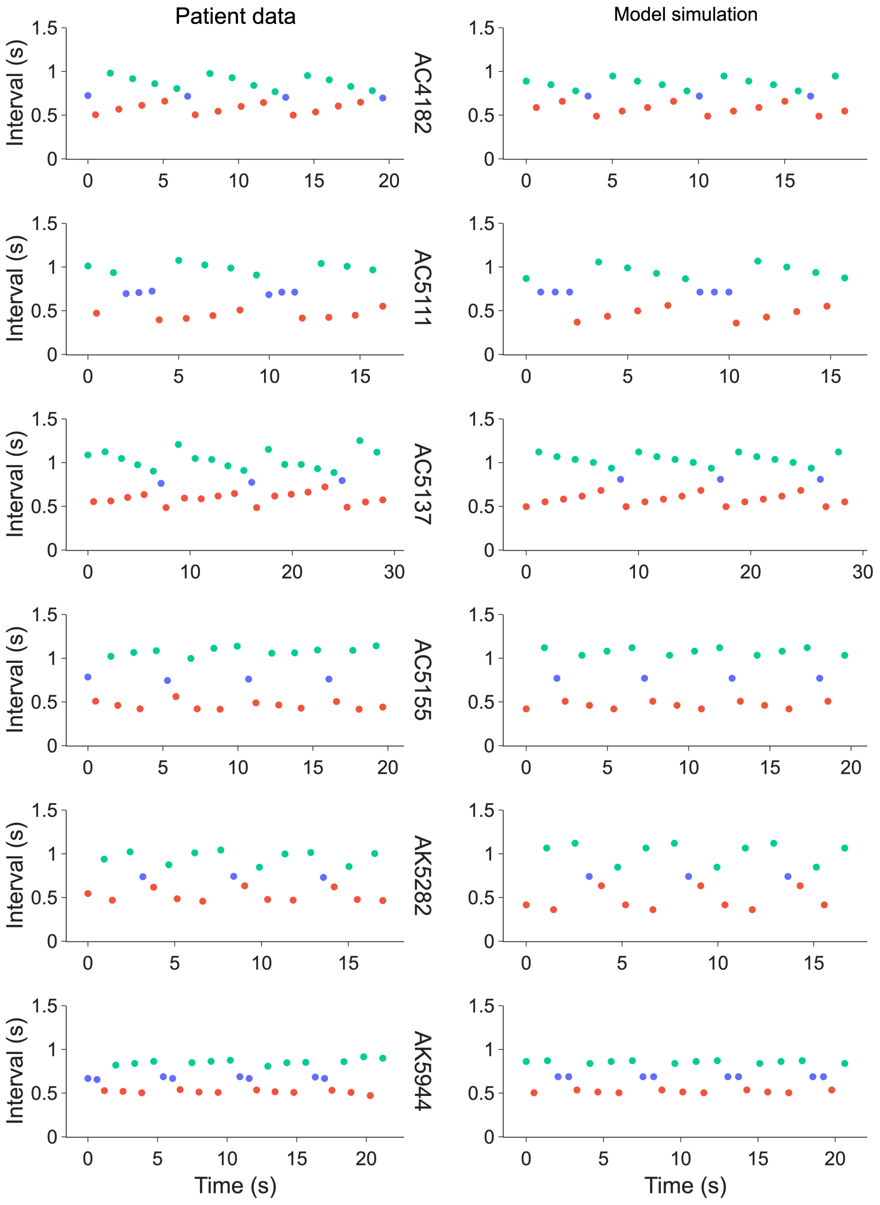


**Figure S3**: **Sections of patient data with cycling coupling intervals (left) and corresponding model simulation (right).** Each panel shows inter-beat intervals from six different patient records identified as exhibiting a cycling coupling interval. Intervals are color-coded as follows: blue for sinus–sinus, red for sinus–ectopic, green for ectopic–sinus, and purple for ectopic–ectopic beats. These sections are used to fit the model parameters $t_e$, $t_{\mathrm{lag}}$, and $\phi_r$ for each record by minimizing the mean absolute error between the model simulation and the data.

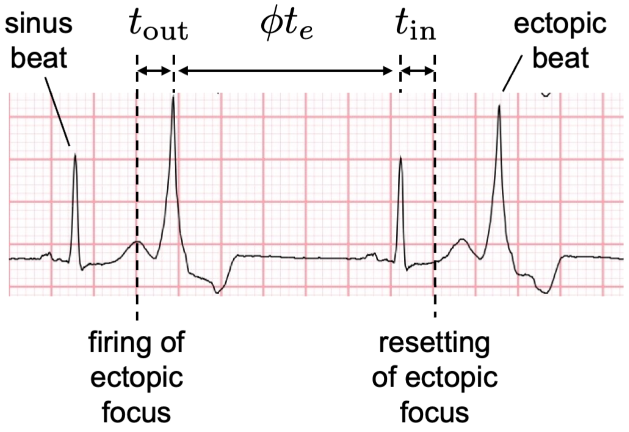


**Figure S4**: **Illustration of conduction time into and out of the ectopic focus.** This ECG shows a bigeminal rhythm (alternating sinus beats and ectopic beats). Assuming the ectopic beats originate from an ectopic pacemaker, we define the phase $\phi$ as the time between the ectopic beat and the subsequent sinus beat divided by the natural period of the ectopic pacemaker $t_e$. We assume that there is a conduction time $t_{\text{out}}$ from the firing of the ectopic pacemaker to the time of the PVC. We also assume there is a conduction time $t_{\text{in}}$ from the time of the sinus beat to the modulation of the ectopic focus. The shifted phase $\phi + t_{\text{lag}}/t_{\text{e}}$ therefore represents the phase at which sinus stimulus arrives at the ectopic focus.

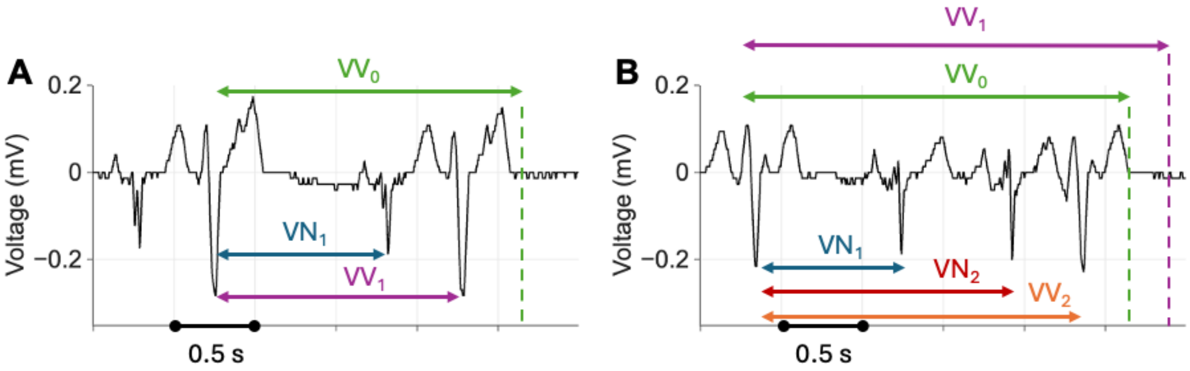


**Figure S5**: Notation for beat-to-beat intervals. (A) Trace shows a section of ECG from record AC5137 during bigeminy (one sinus beat in between ectopic beats). $VN_1$ denotes the interval from the ectopic beat to the (first) sinus beat. $VV_0$ denotes the intrinsic ectopic cycle length. $VV_1$ denotes modified ectopic cycle length due to the sinus beat. (B) Trace shows a section of the same record in trigeminy (two sinus beats in between ectopic beats). $VN_1$ denotes the interval from the ectopic beat to the first sinus beat. $VN_2$ denotes the interval from the ectopic beat to the second sinus beat. $VV_0$ denotes the intrinsic ectopic cycle length. $VV_1$ denotes modified ectopic cycle length due to the first sinus beat. $VV_2$ denotes the modified cycle length due to the first and second sinus beats combined.

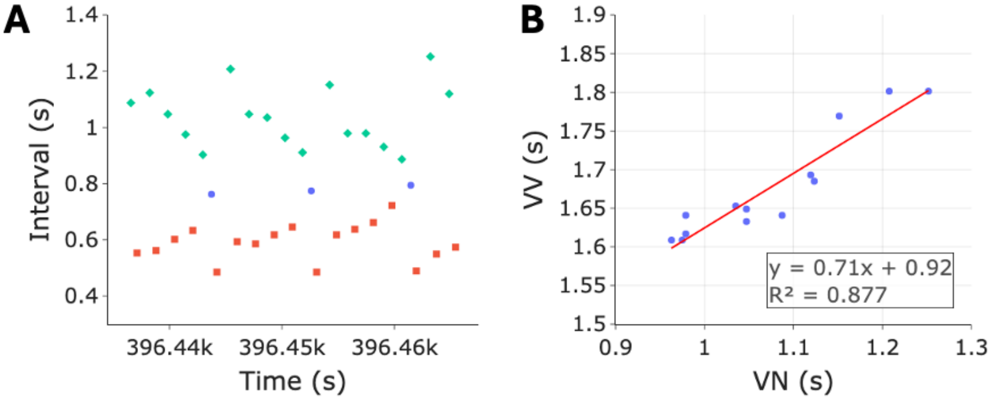


**Figure S6**: **Estimating the PRC resetting slope (S) from the ECG.** (A) Inter-beat intervals for a 30 s section of ECG from record AC5137. Intervals are either between two sinus beats (blue), between a sinus and an ectopic beat (red) or between an ectopic and sinus beat (green). (B) The interval between two consecutive ectopic beats (VV) as a function of the interval between the first ectopic and intervening sinus beat (VN) during periods of NIB=1 (a single sinus beat between two ectopic beats). The slope of the linear regression through the points provides an estimation for the slope of the resetting curve (S).

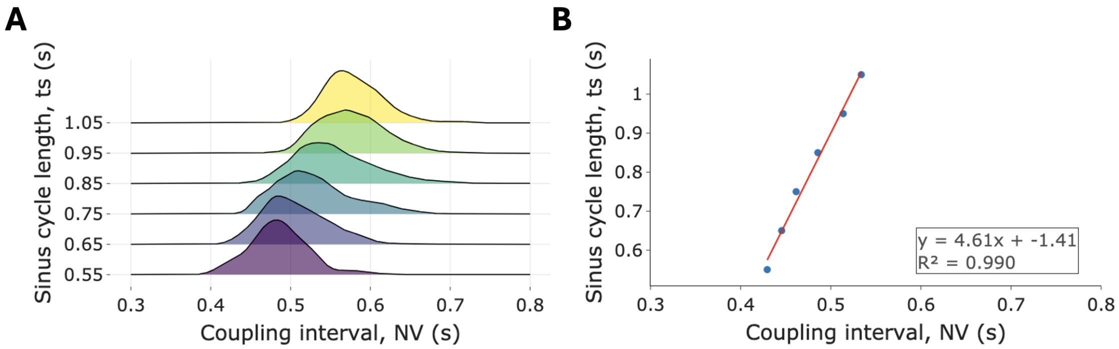


**Figure S7**: **Estimating the refractory period ($\theta$) from the ECG**. (A) Distributions of the coupling interval (NV) at different values of the sinus cycle length (bins of width 0.1 s) for record AC5137. (B) Linear regression through the points that mark the 5th percentile of each distribution. The refractory period at a given sinus cycle length is taken as the coupling interval on this linear regression minus a fixed amount $\epsilon = 0.05$ s. Let $m$ and $c$ be the slope and intercept of the linear regression, then $\theta(t_s) = (t_s - c)/m - \epsilon$.

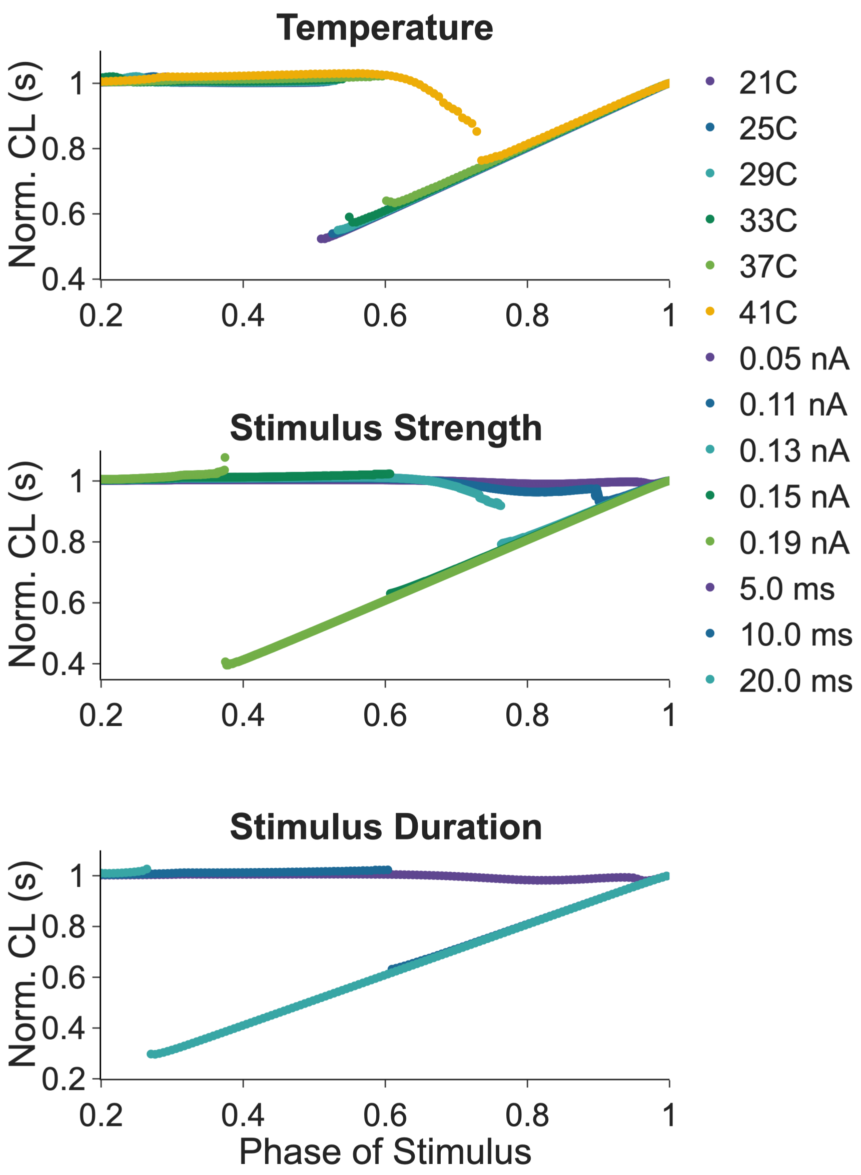


**Figure S8**: **Effects of temperature, stimulus strength, and duration on the PRC from the Paci et al. 2020 ionic model** (**A**) Stimulus strength = 0.15 nA, stimulus duration = 10 ms, temperature from 21°C in indigo to 41°C in yellow. (**B**) stimulus strength from 0.05 nA in indigo to 0.19 nA in green, stimulus duration = 10 ms, temperature = 37°C. (**C**) stimulus strength = 0.15 nA, stimulus duration from 5 ms in indigo to 20 ms in blue, temperature = 37°C.

| Aggregate | $A$ | $\phi_r$ | $B$ | $S$ | RMSE |
|---|---|---|---|---|---|
| A | 60 | 0.71 | 50 | 0.80 | 0.056 |
| B | 40 | 0.57 | 10 | 0.93 | 0.082 |
| C | 120 | 0.57 | 0.73 | 0.80 | 0.124 |
| D | 20 | 0.42 | 10 | 0.93 | 0.155 |
| E | 25 | 0.40 | 10 | 0.95 | 0.103 |
| F | 30 | 0.38 | 20 | 0.95 | 0.058 |

**Table S1**: Optimal parameters for fitting the PRC function (Eqn. (2)) to the experimental PRCs shown in Fig. S1. Aggregate labels correspond to the panel labels in Fig. S1. The root mean squared error (RMSE) is used to assess the goodness of fit.

| Label | Definition | Units | Range/Value | Estimation method |
|---|---|---|---|---|
| $t_s$ | sinus cycle length | s | 0.36-1.61 | Average of RR intervals within a 30 s window |
| $t_e$ | ectopic cycle length | s | 1.50-2.20 | Fit to ECG data (Fig. S3) |
| $t_{\text{lag}}$ | time between sinus beat and resetting of ectopic focus | s | 0.35-0.48 | Fit to ECG data (Fig. S3) |
| $\theta$ | refractory period | s | 0.25-0.68 | Lower bound of NV intervals at each heart rate (Fig. S7) |
| $S$ | PRC resetting slope | – | 0.05-0.95 | Slope of linear regression of VV-VN plot (Fig. S6) |
| $\phi_r$ | PRC discontinuity | – | 0.40-0.80 | Fit to ECG data (Fig. S3) |
| $m$ | PRC delay quotient | – | 45 | Obtained from experimental data |
| $p$ | PRC delay exponent | – | 2.5 | Obtained from experimental data |
| $r$ | PRC delay asymptote | – | 0.745 | Obtained from experimental data |
| $N$ | PRC hill coefficient | – | 35 | Obtained from experimental data |

**Table S2**: Parameters, definitions, and estimation methods for the modulated parasystole model. Ranges/values are across the 7 patients identified as having cycling coupling intervals given in Table S3. PRC: phase response curve; s: seconds; –: dimensionless.

| Record ID | $t_s$ (s) | $\theta$ (s) | $t_e$ (s) | $t_{\text{lag}}$ (s) | $S$ | $\phi_r$ | Prop. record fit (± 5%/10%) |
|---|---|---|---|---|---|---|---|
| AC4182 | 0.46-1.10 | 0.40-0.48 | 1.88 | 0.44 | 0.80 | 0.66 | 75.5% / 99.7% |
| AC5111 | 0.36-1.29 | 0.25-0.45 | 1.51 | 0.25 | 0.08 | 0.75 | 63.4% / 100.0% |
| AC5137 | 0.50-1.28 | 0.35-0.47 | 2.00 | 0.46 | 0.80 | 0.71 | 72.2% / 100.0% |
| AC5155 | 0.42-1.30 | 0.33-0.41 | 1.50 | 0.46 | 0.93 | 0.48 | 73.2% / 95.3% |
| AK5282 | 0.52-1.14 | 0.40-0.50 | 1.61 | 0.30 | 0.75 | 0.41 | 95.0% / 99.8% |
| AK5942 | 0.73-1.61 | 0.64-0.68 | 2.20 | 0.48 | 0.60 | 0.40 | 81.3% / 92.3% |
| AK5944 | 0.53-1.27 | 0.42-0.52 | 1.50 | 0.41 | 0.59 | 0.40 | 63.4% / 89.3% |

**Table S3**: Model parameter values determined for each patient, along with the proportion of 30-second segments of the ECG record that are well described by the model within a ±5% and ±10% variation in parameter values. The sinus cycle length ($t_s$) is measured for each 30-second section of the record and given as a range. The refractory period ($\theta$) is determined from a linear function of $t_s$, derived from the distribution of coupling intervals across heart rates (Fig. S7). The ectopic cycle length ($t_e$), the lag time ($t_{\text{lag}}$), and the PRC discontinuity ($\phi_r$) are determined by fitting the model to a 30-second section identified as having cycling coupling intervals. The slope of the PRC ($S$) is determined by the slope in the plot of VV intervals against VN intervals (Fig. S6). The model is considered a good fit if the Kolmogorov–Smirnov test does not detect a statistically significant difference ($p > 0.05$) between the interbeat interval distributions of the patient and the model.

### References and Notes


1. Z. Jia, *et al.*, Stimulating cardiac muscle by light: cardiac optogenetics by cell delivery. *Circulation: Arrhythmia and Electrophysiology* **4** (5), 753–760 (2011).

2. Y.-L. Huang, A. S. Walker, E. W. Miller, A photostable silicon rhodamine platform for optical voltage sensing. *Journal of the American Chemical Society* **137** (33), 10767–10776 (2015).

3. A. Klimas, G. Ortiz, S. C. Boggess, E. W. Miller, E. Entcheva, Multimodal on-axis platform for all-optical electrophysiology with near-infrared probes in human stem-cell-derived cardiomyocytes. *Progress in biophysics and molecular biology* **154**, 62–70 (2020).

4. M. R. Guevara, A. Shrier, L. Glass, Phase resetting of spontaneously beating embryonic ventricular heart cell aggregates. *American Journal of Physiology-Heart and Circulatory Physiology* **251** (6), H1298–H1305 (1986).

5. M. Courtemanche, L. Glass, M. D. Rosengarten, A. Goldberger, Beyond pure parasystole: promises and problems in modeling complex arrhythmias. *American Journal of Physiology-Heart and Circulatory Physiology* **257** (2), H693–H706 (1989).

6. F. Santoro, *et al.*, Ventricular fibrillation triggered by PVCs from papillary muscles: clinical features and ablation. *Journal of cardiovascular electrophysiology* **25** (11), 1158–1164 (2014).

7. T. Bury, *et al.*, The inverse problem for cardiac arrhythmias. *Chaos: An Interdisciplinary Journal of Nonlinear Science* **33** (12) (2023).

8. K. Takayanagi, *et al.*, Ectopic cycle length estimation from the quantified distribution patterns of ventricular bigeminy and trigeminy. *Heart Rhythm O2* **2** (2), 138–148 (2021).

9. M. Paci, *et al.*, All-optical electrophysiology refines populations of in silico human iPSC-CMs for drug evaluation. *Biophysical journal* **118** (10), 2596–2611 (2020).
